# Dynamic changes in structure and function of brain mural cells around chronically implanted microelectrodes

**DOI:** 10.1101/2024.06.11.598494

**Authors:** Steven M. Wellman, Adam M. Forrest, Madeline M. Douglas, Ashwat Subbaraman, Guangfeng Zhang, Takashi D.Y. Kozai

## Abstract

Integration of neural interfaces with minimal tissue disruption in the brain is ideal to develop robust tools that can address essential neuroscience questions and combat neurological disorders. However, implantation of intracortical devices provokes severe tissue inflammation within the brain, which requires a high metabolic demand to support a complex series of cellular events mediating tissue degeneration and wound healing. Pericytes, peri-vascular cells involved in blood-brain barrier maintenance, vascular permeability, waste clearance, and angiogenesis, have recently been implicated as potential perpetuators of neurodegeneration in brain injury and disease. While the intimate relationship between pericytes and the cortical microvasculature have been explored in other disease states, their behavior following microelectrode implantation, which is responsible for direct blood vessel disruption and dysfunction, is currently unknown. Using two-photon microscopy we observed dynamic changes in the structure and function of pericytes during implantation of a microelectrode array over a 4-week implantation period. Pericytes respond to electrode insertion through transient increases in intracellular calcium and underlying constriction of capillary vessels. Within days following the initial insertion, we observed an influx of new, proliferating pericytes which contribute to new blood vessel formation. Additionally, we discovered a potentially novel population of reactive immune cells in close proximity to the electrode-tissue interface actively engaging in encapsulation of the microelectrode array. Finally, we determined that intracellular pericyte calcium can be modulated by intracortical microstimulation in an amplitude- and frequency-dependent manner. This study provides a new perspective on the complex biological sequelae occurring the electrode-tissue interface and will foster new avenues of potential research consideration and lead to development of more advanced therapeutic interventions towards improving the biocompatibility of neural electrode technology.

## 2.0 INTRODUCTION

Advances in implantable microelectrode technology are evolving at a rapid rate, with recent electrode designs offering the capability to interface with hundreds or thousands of neurons at a time [1-4]. Recording or manipulating brain activity with high spatial and temporal resolution has revolutionized medical treatment for individuals who suffer from neurological disabilities or debilitating brain disease [5, 6]. Despite this progress, there remains challenges using neural microelectrodes for long-term clinical applications due to unpredictable immune responses to an indwelling foreign body [7-12]. Investigators have spent the last three decades attempting to discern the biological origins of a gradual neurodegenerative fate surrounding chronic microelectrodes, yet efforts to identify all of the contributors to the brain’s foreign body response to implanted devices have been confounded by unexplained variability in research findings [13-15]. Realizing the potential of a fully integrated, long-lasting neural electrode interface necessitates a more in-depth understanding of this experimental variability in the foreign body response to neural microelectrode implants.

One theory of biological variability to neural microelectrodes involves the currently unknown consequence of neurovascular damage to device implantation [7]. Insertion of large, mechanically non-compliant electrode arrays often with complex, three-dimensional geometries unavoidably result in damage to underlying vasculature. Sharp, penetrating electrodes can produce variable amounts of bleeding within the brain both near and far from the site of device implantation depending on insertion conditions, such as when implanting through larger descending blood vessels versus directly into the capillary bed [16]. Implantation of these devices can sever or rupture nearby blood vessels leading to influx of plasma proteins that are normally kept out of the brain by the blood-brain barrier [17]. The sudden influx of harmful plasma proteins into the parenchyma initiates a cascade of cellular and tissue processes involving neuroinflammation, glial activation, neurodegeneration, and potentially pericyte dysfunctions around chronically inserted microelectrodes [12, 16]. Pericytes are mural cells which exist on the abluminal side of the blood-brain barrier and are responsible for various regulatory, homeostatic, and neuroimmune functions within the brain [18]. Situated between neuronal synapses and astrocytic endfeet on one side and the endothelial barrier on the other, pericytes are uniquely positioned to communicate signals between neurons and blood vessels to drive neurovascular coupling within the brain [19]. Despite their significance in supporting the metabolic and waste clearance demands of neurons within the brain, pericytes are currently severely underacknowledged in the study of neural electrode biocompatibility [11, 20].

Immediately following a major insult to the brain, pericytes constrict capillary vessels, reduce blood flow, and subsequently perish [21-23]. This loss in pericyte number is followed by a resurgence of new cells which initiate angiogenesis and re-vascularize damaged tissue [24, 25]. Coverage of cortical blood vessels is an important pericyte function for regulating blood-brain barrier permeability and has been linked to the neuropathology of degenerative brain diseases, such as stroke and Alzheimer’s disease [26]. Due to the ability to regulate vascular permeability, pericytes also contribute to the recruitment of peripheral immune cells following injury [27]. Finally, pericytes reportedly possess mesenchymal stem cell like activity, such as differentiating into microglia-like cells during pathological conditions such as stroke [27, 28]. Nevertheless, pericytes are increasingly emerging as significant perpetuators of the neurodegenerative outcomes in brain injury and disease.

Previously, we demonstrated that pericyte densities fluctuate spatiotemporally around chronically implanted microelectrodes [12]. Until now, the pericyte contribution to the foreign body response following neural probe implantation has been examined solely through the lens of post-mortem tissue sections [12, 20], providing only a snapshot of the tissue response outcomes at the endpoint of the experiment. Two-photon microscopy is an emerging imaging technique to longitudinally visualize the morphology and activity of different cellular populations within the brain in real-time and offers a more detailed characterization of the dynamic tissue response to chronic brain implants [29-33]. Using intravital imaging, we observe dynamic changes of brain pericytes over a 28-day period around chronically implanted microelectrodes. Specifically, we determine that pericytes respond immediately (<30 min) to electrode insertion by constriction of microvessels, and whose Ca^2+^-mediated contraction can be modulated using intracortical microstimulation, followed by gradual remodeling of local vasculature over the course of days to weeks following implantation. Using dual-transgenic mouse models, we also reveal for the first-time a novel reactive immune cell population at the device-tissue interface. In summary, our work highlights a crucially understudied component of the foreign body response to intracortical electrodes and the novel findings presented here will ultimately benefit the future development of more advanced intervention strategies and innovative designs of neural implant technology.

## 3.0. METHODS

### 3.1. Experimental animal models

Tg(Cspg4-DsRed.T1)1Akik mice (male, 22-30 g, 8-12 weeks old) expressing fluorescent DsRed protein in vascular smooth muscle cells (vSMCs) and pericytes were obtained from Jackson Laboratories (Stock #008241, Bar Harbor, ME). Dual-fluorescent pericyte and microglia models were generated by crossing male Tg(Cspg4-DsRed.T1)1Akik mice (Cspg4-DsRed Stock #008241, Jackson Laboratories, Bar Harbor, ME) and female B6.129P-CX3cr1/J mice (CX3cr1-GFP, Stock #005582, Jackson Laboratories, Bar Harbor, ME). To visualize GCaMP expression within pericytes, male B6.Cg-Tg(Pdgfrb-CreERT2)6096Rha/J mice (Pdgfrb-CreERT2, Stock #029684, Jackson Laboratories, Bar Harbor, ME) were crossed with female B6J.Cg-Gt(ROSA)26Sortm96(CAG-GCaMP6s)Hze/MwarJ mice (Ai96(RCL-GCaMP6s), Stock #028866, Jackson Laboratories, Bar Harbor, ME). To induce GCaMP6s expression, PDGFRβ-GCaMP6s mice were injected intraperitoneally with tamoxifen (10 mg/mL in corn oil) at a dose of 100 mg/kg on five consecutive days six weeks prior to surgery. All procedures and experimental protocols were approved by the University of Pittsburgh, Division of Laboratory Animal Resources, and Institutional Animal Care and Use Committee in accordance with the standards for humane animal care as set by the Animal Welfare Act and the National Institutes of Health Guide for the Care and Use of Laboratory Animals.

### 3.2. Surgical probe implantation

Probe implantation surgeries were performed as described previously [29-34]. Prior to surgery, animals were sedated using an anesthetic cocktail mixture of xylazine (7 mg/kg) and ketamine (75 mg/kg). The top of the mouse head was shaved and sterilized with alternating betadine and ethanol scrubs. A single incision was made over the mouse skull followed by removal of excess hair and connective tissue. Adhesive Vetbond was applied to the skull prior to drilling. Two bone screw holes centered over each motor cortex were drilled and fitted with bone screws. A 4x4 mm sized craniotomy was created over both hemispheres. A 4-shank Michigan style microelectrode array was inserted at a 30° angle to a depth of 300 μm below the surface within the right visual cortex (1.5 mm anterior, 1 mm lateral from lambda). The contralateral cortex was left un-implanted as an un-injured control region for comparison. A silicone elastomer (Kwik-sil) was used to fill the space within the craniotomy above the brain prior to covering with a glass coverslip and sealing the edges of the glass with dental cement for optical imaging. A separate cohort of C57BL/6J mice were implanted for post-mortem histological analysis as described previously [12]. Similar implantation procedures were repeated in these mice with the exception of the formation of a drill-sized craniotomy over the visual cortex (1.5 mm anterior, 1 mm lateral from lambda) in which a single shank Michigan style microelectrode array was lowered to a 1.6 mm depth into the cortex at a 90° angle. Kwik-sil was applied to fill the space within the craniotomy and around the electrode prior to sealing within a dental cement headcap. Ketofen (5 mg/kg, Covetrus, Inc., Portland, ME) was provided post-operatively on the day of surgery and up to 2 days post-surgery to all implanted animals.

### 3.3. Two-photon laser-scanning microscopy

Two-photon microscopy was used to track the dynamic pericyte response to chronically implanted microelectrodes. The microscope was equipped with a scan head (Bruker, Madison, WI), a OPO laser (Insight DS+; Spectra-Physics, Menlo Park, CA), non-descanned photomultiplier tubes (Hamamatsu Photonics KK, Hamamatsu, Shizuoka, Japan), and a 16X, 0.8 numerical aperture water-immersive objective lens (Nikon Instruments, Melville, NY). A 920 nm & 980 nm two-photon laser wavelength were used for fluorescence excitation in PDGFRβ-GCaMP6s or Cspg4-DsRed mice, respectively. Blood vessels were visualized using either sulforhodamine 101 (intraperitoneal injection, 1 mg/mL) or a FITC-dextran dye (retro-orbital injection, 2 MDa, 10 mg/mL). Volumetric Z-stack ROIs were acquired from both the ipsilateral and contralateral hemisphere for each imaging session (0, 2, 4, 7, 14, 21, and 28 d post-implantation). Images were collected at ∼5 s frame scan rate (1024×1024 pixels, ∼5.0 µs dwell time, ∼1.5-2× optical zoom) and 2-3 µm step size beginning at the pial surface all the way along the full depth of the electrode implant.

### 3.4. Intracortical microstimulation

Electrical stimulation trials in PDGFRβ-GCaMP6s mice were performed 1 week following device implantation to allow the animal to recover and to assess the impacts of intracortical microstimulation on pericyte function without the interference of residual anesthesia from surgery. Electrical stimulation was administered using a Tucker-Davis Technologies RZ6D system and IZ2 stimulator (Tucker-Davis Technologies, Alachua, FL). Stimulation trials consisted of a charged balanced, bi-phasic, cathodic-leading asymmetric waveform with 200 µs leading phase, 100 µs delay, and 400 µs lagging phase at half-amplitude. Stimulation conditions varied by stimulation amplitude (5, 10, and 15 µA) and frequency (10 Hz burst, 10, 40, and 100 Hz continuous). Stimulation trials were comprised of 30 s baseline, 30 s stimulation duration, and 60 s post-stimulation during two-photon imaging. Pericyte ROIs were imaged at 128 x 128 pixels, 2 µs dwell time, and 4x optical zoom. All stimulation waveforms were delivered between 1-3 nC/phase, which is within the safety limit of electrical stimulation using intracortical microelectrodes. All stimulation amplitude and frequency combinations were delivered in random order and repeated at least three times.

### 3.5. Post-mortem immunohistochemistry

C57BL/6J mice with perpendicularly implanted probes were sacrificed at 1-, 7-, and 28-days (*n* = 5) following insertion for immunohistochemical analysis. Briefly, mice were anesthetized with a xylazine/ketamine cocktail and transcardially perfused with 1X PBS followed by 4% paraformaldehyde (PFA). Probes were left intact attached the skull and within the brain and post-fixed in 4% PFA at 4°C overnight. Then, brains were extracted and allowed to equilibrate in 15% and then 30% sucrose at 4°C for 12-24 hours each. Samples were embedded and frozen in a 2:1 PBS:optimal cutting temperature (OCT) media prior to cryosectioning. Tissue sections of 25 μm thickness were collected horizontally along the entire depth of the implant. Prior to staining, sections were re-hydrated in 1X PBS for 5 min. For antigen retrieval, slides were incubated in 0.01 M sodium citrate buffer for 30 min at 60C and then in peroxidase blocking solution (10% v/v/ methanol and 3% v/v/ hydrogen peroxide) for 20 min on a table shaker at room temperature. Prior to antibody staining, sections were permeabilized with a solution of 1% triton X-100 and 10% donkey serum in PBS for 30 min at RT. To eliminate non-specific binding, sections were then blocked with donkey anti-mouse IgG fragment (fab) or 647 conjugated anti-mouse IgG fragment (Fab) for 2 h at 1:13 or 1:16 dilution at RT. Sections were then washed 8 x 4 min in 1X PBS prior to incubating in a primary antibody solution for 12-18 hr at 4°C. Primary antibodies used were rabbit anti-NG2 (1:200, ab5320, Sigma Aldrich, St. Louis, MO), goat anti-PDGFRβ (1:100, AF1042, R&D Systems, Minneapolis, MN), mouse anti-Ki67 (1:100, 550609, BD Biosciences), and anti-Tomato lectin (1:250, B-1175, Vector Labs, Newark, CA).

The following day, sections were rinsed with 1X PBS three times for 5 minutes each prior to incubation with the following secondary antibodies at 1:500 in 1X PBS for 2 hr at RT: 405 donkey anti-rabbit, 488 donkey anti-goat, and 568 donkey anti-mouse (Sigma Aldrich, St. Louis, MO). After secondary antibody incubation, sections were then rinsed again with 1X PBS 3×5 min. Slides were mounted with Fluoromount-G, coverslipped, and allowed to dry overnight at RT before being stored in 4°C.

### 3.6. Data analyses

#### 3.6.1. Pericyte coverage analysis

Pericyte vascular coverage following chronic electrode implantation was assessed via immune-stained sections for PDGFRβ+ pericytes and lectin+ blood vessels. Both PDGFRβ and lectin stains were binarized by setting a threshold fluorescence intensity value of 2.5 times the standard deviation above the mean. Binarized images were then binned every 50 µm up to 300 µm away from the center of probe implantation. Vascular coverage was measured as the number of overlapping PDGFRβ+ and lectin+ pixels divide by the total number of lectin+ pixels within each bin. Pericyte coverage was then normalized to the contralateral hemisphere.

#### 3.6.2. Quantification of Cspg4+/Cx3cr1+ cell distribution and surface encapsulation

The geometric distribution of the Cspg4+/Cx3cr1+ cell population was determined by generating a three-dimensional distance map of the electrode surface, as described previously [30]. Briefly, the orientation angle of the electrode within the imaged volume was determined using the ‘Interactive Stack Rotation’ plugin in ImageJ (National Institute of Health) and used to rotate the electrode into a single horizontal plane. Then, a binary mask was created by manually defining an outline of the probe. This binary mask was then rotated back into the original electrode orientation and a distance transformation was applied to the resultant 3D mask using the ‘bwdist’ function in Matlab (MathWorks, Boston, MA) to form a distance map. The spatial coordinates of identified cell counts were then referenced using this distance map for each mouse and each time point quantified.

Cspg4+ and CX3cr1+ cellular encapsulation was determined as a percentage of fluorescent signal covering the surface of the implanted probe, as previously described [31-33]. Briefly, each image volume was rotated and resliced using the ImageJ plugin “Interactive Stack Rotation” to orient the entire length of the angled probe within a single 2D plane. Then, a small volume of tissue above the probe was sum slice projected into a single image and a binary mask was created using an “IsoData” thresholding algorithm within ImageJ. The percentage of fluorescent signal was taken as the number of non-zero pixels within a manually defined outline of the probe over the total area measured. The final dimensions of the outlined probe area were verified prior to thresholding for accuracy.

#### 3.6.3. Pericyte GCaMP fluorescence intensity analysis

For acute insertion trials, *PDGFRβ-GCaMP6s* mice were imaged for up to 20 min post-insertion. ROIs of pericyte soma and processes were manually defined using the ROI manager in ImageJ. Fluorescence intensities were calculated as ΔF/F_0_, where F_0_ is the average baseline fluorescence intensity averaged over 1 min pre-insertion. Activated pericytes were defined as cells whose ΔF/F_0_ was greater than F_0_ plus three times standard deviation (3*STD) of baseline recordings. Fluorescence intensities were normalized to pre-insertion recording of calcium activity. For electrical stimulation trials, ROIs of pericyte soma were defined manually using ImageJ, same as above. Fluorescent calcium traces were extracted using a built-in image processing toolbox, Cellular and Hemodynamic Image Processing Suite (CHIPS) [35], based on MATLAB (R2017b, Mathworks). Vessel diameter traces were collected by manually defining cross-sectional lines across the vessel lumen and using VasoMetrics, a custom ImageJ macro [36]. Fluorescent calcium or vessel diameter measurements were normalized using the average signal value during 30 s pre-stimulation period as baseline. Mean ΔF/F_0_ or ΔD/D_0_ was calculated as the average change in normalized calcium fluorescence or vessel diameter, respectively, during the stimulation period. Duration was calculated as the time required for calcium fluorescence of vessel diameter to return back to baseline following stimulation onset. Area under or over the curve was calculated as the definite integral of the normalized calcium fluorescence or vessel diameter, respectively, during the stimulation period.

### 3.7. Statistics

A one-way or two-way ANOVA (*p* < 0.05) was used to assess for differences in cell counts, coverage, motility over time, distance away from the electrode, and pericyte calcium response to intracortical microstimulation. Significant pairwise differences were assessed post hoc using a Tukey’s Honest Significant Difference (HSD) test.

## 4.0. RESULTS

In order to reveal potential alterations in the structure and function of brain pericytes following neural electrode implantation, we utilized two-photon microscopy to longitudinally assess dynamic changes in pericyte fate and morphology around chronically implanted microelectrode arrays. The *Cspg4-DsRed* model was chosen to identify and characterize brain pericytes based on differences in their structure, morphology, and anatomical position within the brain (**Fig. 1**). *Cspg4-DsRed* mice were implanted with a four-shank Michigan style microelectrode under an optical window for longitudinal imaging over a 28-day implantation period.

**Figure 1.**
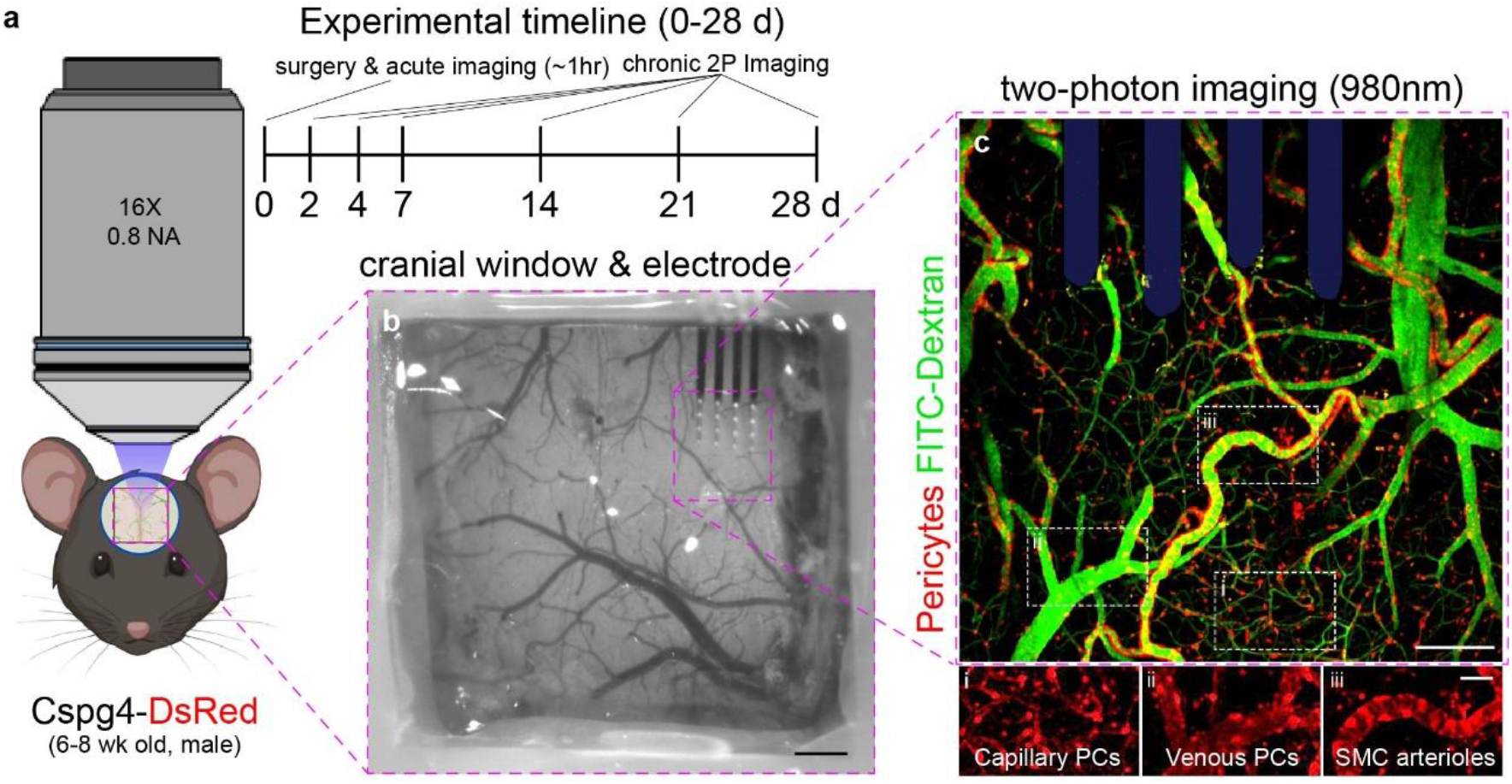
Experimental setup to visualize dynamic changes of vascular mural cells around chronically implanted microelectrodes. (a) Schematic of two-photon visualization of brain tissue through cranial glass window in Cspg4-DsRed mice over a 28-day implantation period. (b) Surgical image of multi-shank microelectrode implanted in mouse cortex underneath glass cranial window. Scale bar = 1 mm. (c) Vascular pericytes around an implanted multi-shank microelectrode array within Cspg4-DsRed mouse model. The Cspg4-DsRed mouse model reveals alterations in the structure and morphology of different distinct classes of brain mural cells, such as capillary pericytes (i), venous pericytes (ii), and smooth muscle cell (SMC) arteriole pericytes (iii). Scale bar = 100 μm, 25 μm (inset).

### 4.1. Pericytes constrict capillaries and increase in intracellular Ca^2+^ following electrode insertion

Pericytes possess contractile proteins that allow them to regulate cerebral blood flow by exerting mechanical forces on blood vessels [36, 37]. Perturbations to the brain can produce contractions of pericyte somata and constriction of microvessels, often time to a pathological extent [38, 39]. To determine whether pericytes respond in a similar manner to electrode insertion, we used intravital imaging to capture changes in pericytes in real-time around an implanted microelectrode (**Fig. 2a**). We demonstrate that pericytes are present at positions along the capillary network where occlusions in blood flow and constriction of microvessels are apparent (**Fig. 2b**). For capillary constriction, we show that vessels are constricted beyond baseline diameter at the position of pericyte soma (**Fig. 2c**). We further confirm pericyte-mediated constriction of microvessels by staining explanted brain tissue using PDGFRβ for pericytes and tomato lectin for the vasculature at 7 days post-implantation in wild-type mice (**Fig. 2d**). Intracellular calcium ([Ca^2+^]) levels regulate the contractile machinery necessary for pericytes to mechanically constrict around blood vessels [36, 37]. To determine whether electrode implantation alters pericyte Ca^2+^ activity, we implanted an electrode into the brains of *PDGFRβ-GCaMP6s* mice express GCaMP within PDGFRβ+ pericyte cells (**Fig. 2e**). There was a transient increase in Ca^2+^ within both pericyte soma and processes in the initial minutes following insertion which returned back to baseline levels after 20 min following implantation. While the temporal pattern of Ca^2+^ changes were similar between both soma and processes over the course of electrode insertion, we determined that pericyte soma demonstrated significantly larger increases in normalized fluorescence changes from baseline compared to pericyte processes (**Fig. 2f**, *p* < 0.001, two-way ANOVA with Tukey’s HSD test). Overall, we demonstrate that pericytes produce deformations on cortical microvessels around implanted microelectrodes and respond to electrode insertion through transient increases intracellular calcium.

**Figure 2.**
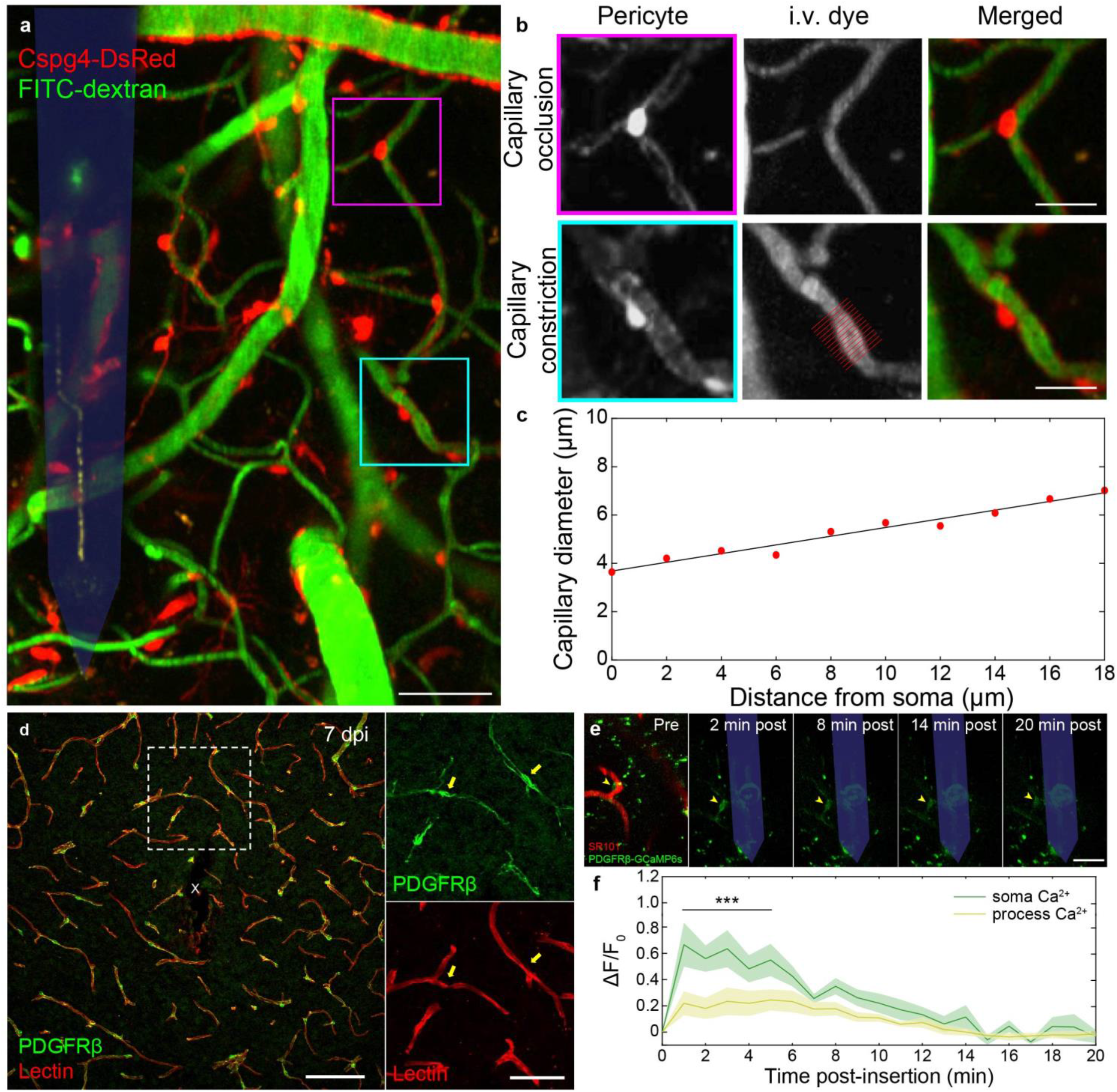
Pericytes constrict capillary vessels and transiently increase in intracellular Ca^2+^ following electrode insertion. (a) Two-photon image of pericytes (Cspg4-DsRed, *red*) and blood vessels (FITC-dextran, *green*) around an implanted microelectrode (*shaded blue)*. Scale bar = 50 μm. (b) Inset of (a) demonstrating regions of capillary occlusion and constriction overlap with position of pericyte soma. Scale bar = 20 μm. (c) Quantification of change in capillary diameter of with respect to distance from center of cell soma. (d) Histological image demonstrating pericytes (PDGFRβ, *green*) located on regions of deformed capillary (Lectin, *red*) structures at 7 days post-implantation of a microelectrode array (*white ‘x’)*. Scale bar = 50 μm, 25 μm (inset). (e) Time course of a transient increase in intracellular pericyte Ca^2+^ (PDGFRβ-GCaMP6s, *yellow arrowhead*) near an inserted probe over a 20 min implantation period. Scale bar = 50 μm. (f) Normalized fluorescence intensity profile of soma and process compartments of pericyte cell in (e) over 20-minute post-insertion period (*n* = 4 soma and 4 processes, *N* = 2 mice). *** *p* < 0.001. All data is reported as mean ± SEM.

### 4.2. Microelectrode implantation promotes pericyte proliferation and angiogenesis

The acute pericyte response to severe brain injury is followed by a resurgence of proliferating, angiogenic pericytes into the damaged tissues regions. To understand whether pericytes respond similarly to electrode implantation injury, we implanted microelectrode arrays into the brains of *Cspg4-DsRed* mice and monitored the chronic response of pericytes over a 28-day implantation period. Previously, we demonstrated that reactive glial populations proliferate around chronically implanted microelectrodes due to their elevated expression of the proliferative marker, Ki67 [12]. Using immunohistochemistry, we show that pericytes within the microenvironment of implanted electrodes during this period of dynamic tissue remodeling, identified by co-expression of NG2 and PDGFRβ markers, also express Ki67 at 1, 7, and 28 days post-implantation (**Fig. 3a-c**). Ki67+ pericytes were not observed on the contralateral hemisphere (**Fig. 3d-f**), supporting our findings that pericytes respond and proliferate specifically in response electrode implantation injury. The density of co-localized PDGFRβ+ and Ki67+ cells was significantly increased at 50-100 μm from the probe surface between 1- and 7-days post-implantation (**Fig. 3g**, *p* < 0.05 at 50-100 μm and *p* < 0.01 at 100-150 μm, two-way ANOVA with Tukey’s HSD test). While the density of PDGFRβ+/Ki67+ cells was reduced from 7 to 28 days in close proximity to the electrode, densities were significantly increased further away from the site of implantation at 150-200 μm between 1- and 28-days post-insertion (*p* < 0.01, two-way ANOVA with Tukey’s HSD test) and at 200-300 μm at 28 days post-insertion compared to 1 and 7 days post-insertion (*p* < 0.001, two-way ANOVA with Tukey’s HSD test). In support of our post-mortem findings, two-photon imaging experiments revealed an influx of new pericytes accumulating at the electrode-tissue interface starting within the first week up until 28 days post-implantation (**Fig. 3h**). These cells exhibited dynamic cell motility, migrating preferentially toward the surface of the electrode in order to facilitate angiogenesis. The vessels previously occupied by these cells were then devoid of pericyte coverage. Coincidentally, the diameters of these vessels which lack pericyte coverage appear increased, most likely due to the loss of pericyte contact. Overall, these results support the notion that chronic electrode implantation promotes dynamic remodeling and angiogenesis with the local tissue microenvironment through a population of proliferating pericyte cells.

**Figure 3.**
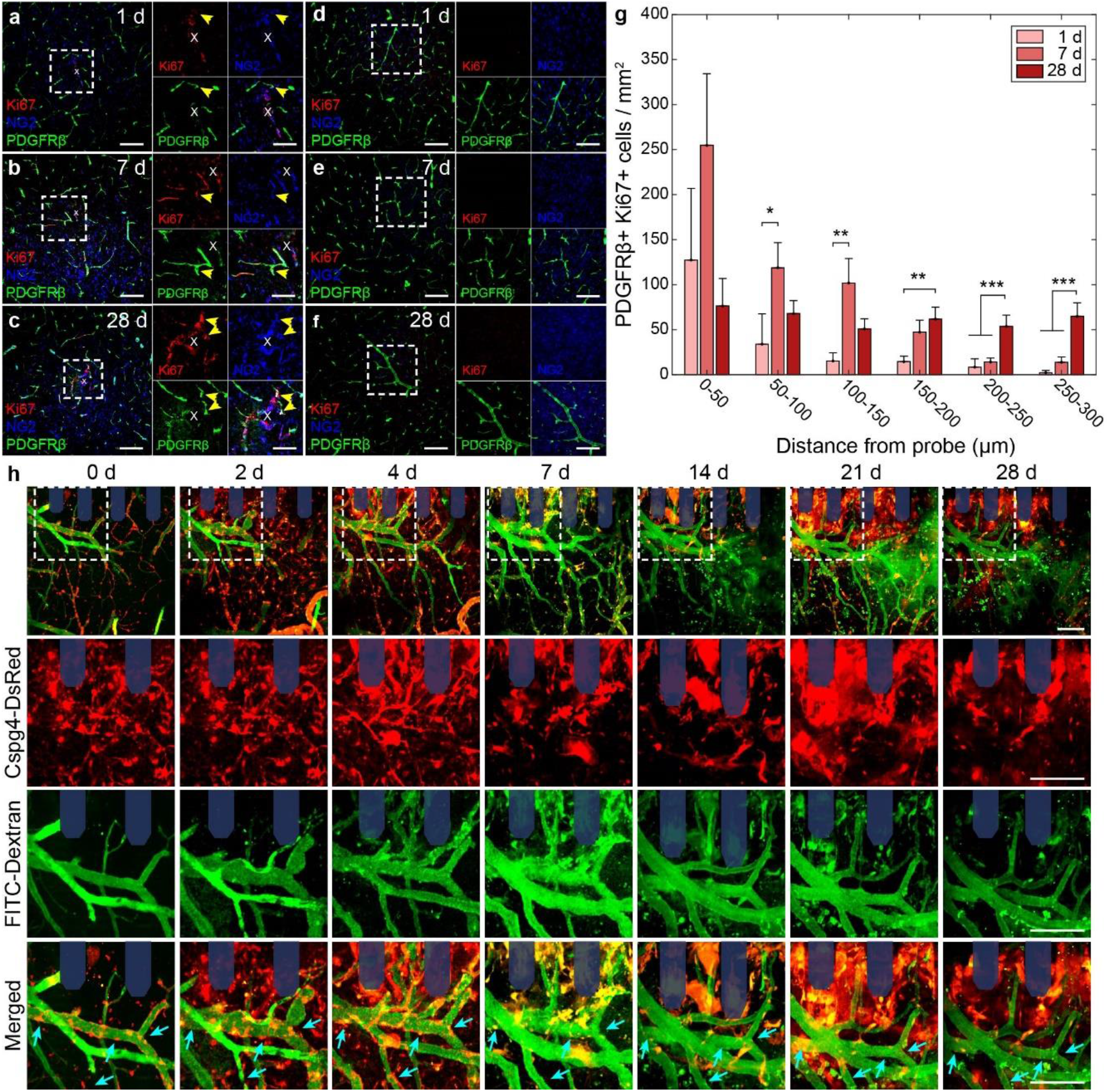
Influx of proliferating pericytes facilitate angiogenesis around implanted microelectrodes. Histological stain of NG2+ (*blue*) and PDGFRβ+ (*green*) pericytes co-expressing proliferative marker Ki67 (*red)* around (a-c) implanted microelectrode (*white ‘x’*) and (d-f) contralateral, unimplanted tissue at 1, 7, and 28 days post-insertion. Yellow arrows point to areas of overlap. Scale bar = 100 μm, 50 μm (inset). (g) Density of PDGFRβ+ and Ki67+ cells in 50 μm bins up to 300 μm at 1, 7, and 28 days post-insertion (*N* = 5 mice). (h) Two-photon representation of time course of dynamic pericyte activity around chronically implanted microelectrodes (*shaded blue*). Cyan arrows indicate areas of in which blood vessels lose pericyte coverage over time. Scale bars = 100 μm. * *p* < 0.05, ** *p* < 0.01, *** *p* < 0.001. All data is reported as mean ± SEM.

### 4.3. Pericyte coverage of blood vessels is impacted by chronic microelectrode implantation

Spatial cell coverage over blood vessels within the brain is an important regulatory function unique to pericytes which allow them to maintain microvascular tone as well as modulate the level of blood-brain barrier permeability [40, 41]. To determine whether pericyte coverage is affected by chronic electrode implantation, we visualized the amount of area overlap between PDGFRβ+ pericytes and lectin+ blood vessels at 1-, 7-, and 28-days post-implantation in post-mortem histological brain sections (**Fig. 4a**). Quantification of normalized area occupied by either PDGFRβ+ pericytes or lectin+ blood vessels was not significantly impacted over a 300 μm radius around the implanted microelectrode at either 1-, 7-, or 28-days post-insertion (**Fig. 4b-c**, *p* > 0.05, two-way ANOVA with Tukey’s HSD test). We did however observe both a reduction in PDGFRβ+ area and an increase in lectin+ area specifically within 0-50 μm from the probe surface at 7 days post-insertion. While statistical analyses of these changes in area alone did not reveal a significant effect, quantification of pericyte coverage of blood vessels (PDGFRβ+/lectin+ area over total lectin+ area) around our implanted microelectrodes resulted in a slight significant reduction in coverage compared to contralateral tissue within a 0-50 μm distance from the probe surface at 7 days post-implantation (**Fig. 4d**, *p* = 0.05, two-way ANOVA with Tukey’s HSD test). These results are in line with our earlier discussed two-photon findings and suggest that pericyte coverage of blood vessels in proximity to the electrode surface may be impacted by the injury sustained during implantation.

**Figure 4.**
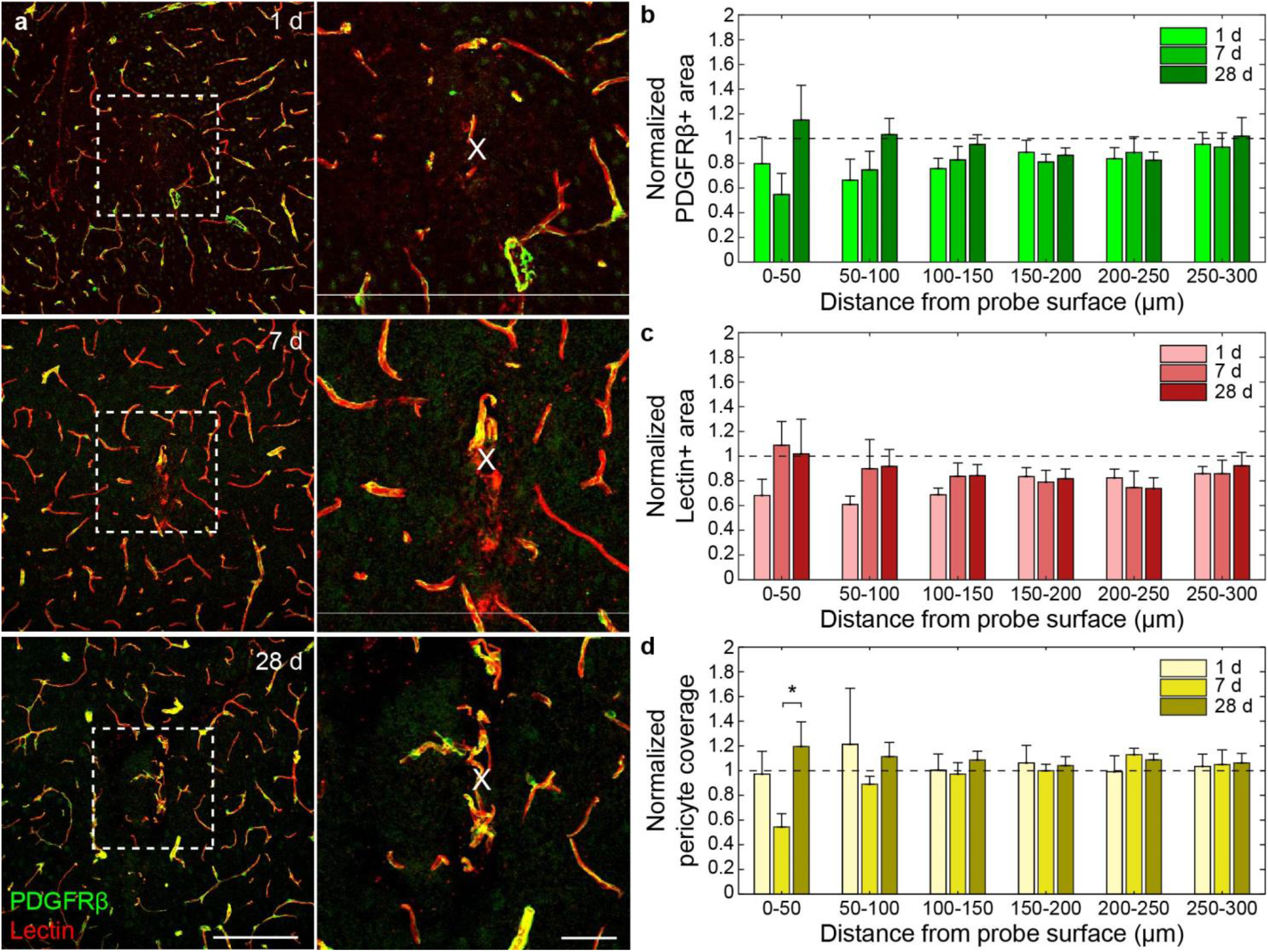
Microelectrode implantation impact on pericyte coverage of blood vessels. (a) Histological representation of pericyte (PDGFRβ, *green*) coverage of blood vessels (Lectin, *red*) around chronically implanted microelectrodes. Yellow represents the overlap between PDGFRβ and lectin stains. Scale bar = 100 μm, 25 μm (inset). (b) Normalized PDGFRβ+ area within 50 μm bins up to 300 μm at 1, 7, and 28 days post-insertion. (c) Normalized lectin+ area within 50 μm bins up to 300 μm at 1, 7, and 28 days post-insertion. (d) Normalized pericyte coverage (PDGFRβ+ area over lectin+ area) reveals significant reduction within 0-50 μm at 7 days post-insertion. *N* = 5 mice. * *p* < 0.05. All data is reported as mean ± SEM.

### 4.4. Involvement of dual-expressing *Cspg4+/CX3cr1+* cells in surface encapsulation of chronically implanted electrodes

Pericytes have been previously implicated in the formation of a fibrotic scar. We previously reported on a temporally distinct pattern of device encapsulation by NG2 glia and microglia cells. To determine whether Cspg4+ expressing pericytes participated in the encapsulation of implanted microelectrodes, we generated a dual-fluorescent *Cspg4-DsRed::CX3cr1-GFP* mouse model to visualize electrode encapsulation of both brain cell populations within the same animal (**Fig. 5a**). In the healthy, intact brain, both pericytes and microglia occupy spatially distinct tissue regions (**Fig. 5b**). With chronic electrode implantation, we determined that both Cspg4+ and CX3cr1+ cells gradually encapsulate the electrode surface over a 28-day implantation period (**Fig. 5c**). Interestingly, we observed distinct regions on the electrode surface in which DsRed+ and GFP+ cells overlap. Upon quantitative measurement, we showed that the encapsulation region of these dual-labeled GFP+/DsRed+ cells were temporally distinct from that of DsRed+ and GFP+ areas alone (**Fig. 5d**). Initially, coverage of the electrode was dominated solely by the GFP+ only signal, which was significantly increased compared to DsRed+ only and GFP+/DsRed+ signals up to 2 days post-insertion (*p* < 0.001, two-way ANOVA with Tukey’s HSD test). Then, at 4 days post-insertion, there was an increase in combined GFP+/DsRed+ signal compared to both GFP+ and DsRed+ signals alone. After one week, coverage of the electrode surface by DsRed+ only of GFP+/DsRed+ signal gradually declines up to three weeks later, with the GFP+ only signal remains significantly elevated at 28 days post-insertion (*p* < 0.05, two-way ANOVA with Tukey’s HSD test). Whereas surface coverage of individual CX3cr1+ or Cspg4+ cells have been previously recognized, here we report a spatially and temporally distinct pattern of surface encapsulation by dual-labeled CX3cr1+/Cspg4+ cells, suggesting the emergence of a novel reactive brain cell population around chronically implanted microelectrodes.

**Figure 5.**
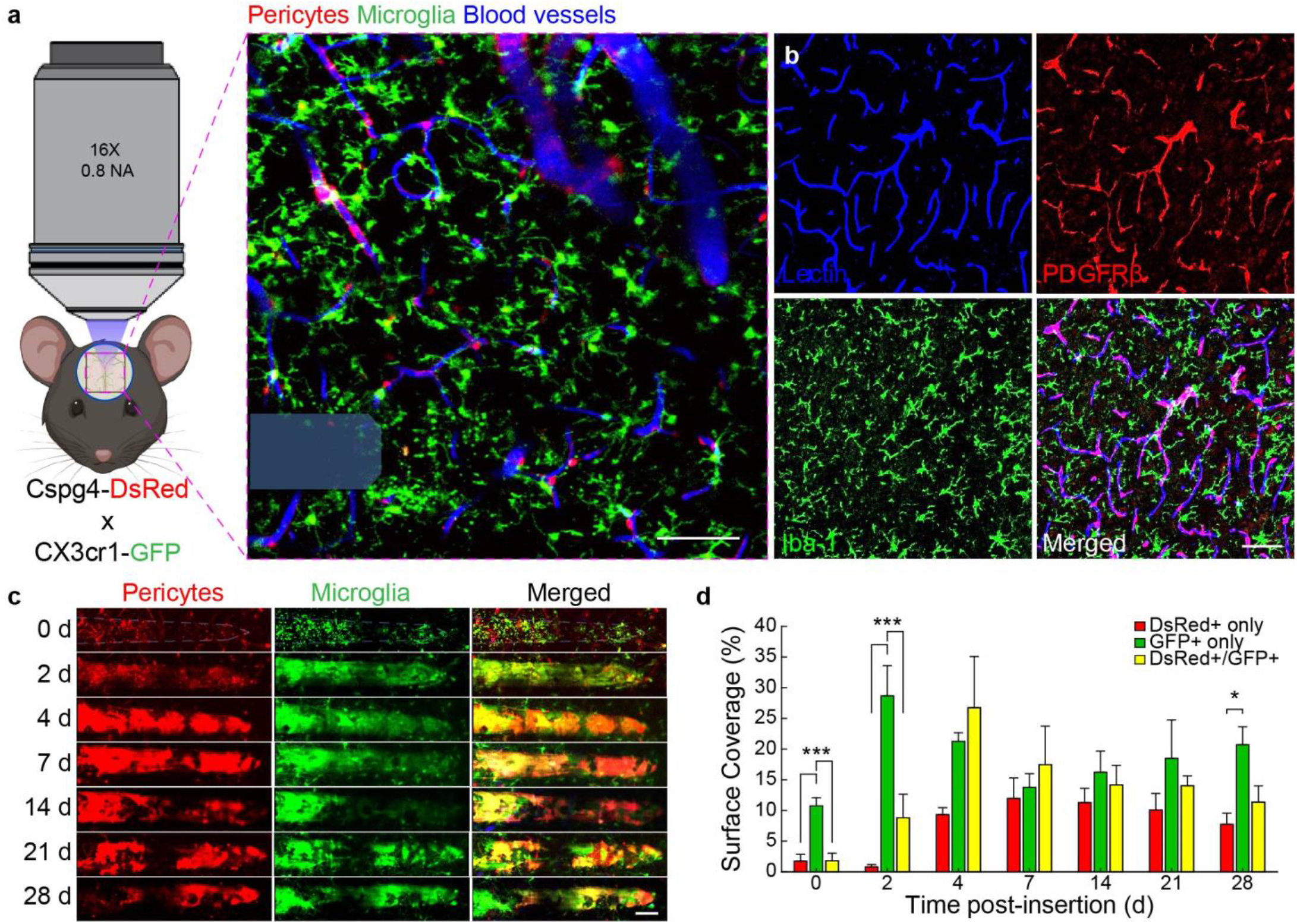
Temporal pattern of microelectrode encapsulation by spatially distinct Cspg4+ and CX3cr1+ cell populations. (a) Two-photon representative image of a microelectrode (*shaded blue*) inserted in a Cspg4-DsRed::CX3cr1-GFP. Blood vessels (*blue*) are visualized using an intravascular fluorescent dye. Scale bar = 50 μm. (b) Histological image of stained blood vessels (*blue*), PDGFRβ+ pericytes (*red*), and Iba-1+ microglia (*green*) in healthy, uninjured brain tissue. Scale bar = 100 μm. (c) Two-photon representative images of surface encapsulation of chronically implanted microelectrodes of Cspg4+ (*red*) and CX3cr1+ (*green*) brain populations. Yellow here denotes overlapping DsRed and GFP fluorescent signals. Scale bar = 50 μm. (d) Percent coverage of DsRed+ only, GFP+ only, and overlapping DsRed+/GFP+ signal over 28-day implantation period (*N* = 4 mice). * *p* < 0.05, *** *p* < 0.001. All data is reported as mean ± SEM.

### 4.5. Novel CX3cr1+/Cspg4+ population of reactive glial cells accumulate at the site of neural electrode implantation

To better understand the spatial relationship between activated microglia and pericyte cells following surface coverage of chronically implanted microelectrodes, we discovered the appearance of a unique population of immune cells at the electrode-tissue interface which simultaneously express GFP+ and DsRed+ fluorophores denoting constitutive expression of both *CX3cr1* and *Cspg4* promoters (**Fig. 6a**). Dual-labeled CX3cr1+/Cspg4+ cells were distinct from individual CX3cr1+ and Cspg4+ cells in morphology and positioning around chronic microelectrodes. Furthermore, these cells were not observed within contralateral (unimplanted) brain regions, suggesting an injury-specific emergence to chronically implanted devices. The appearance of CX3cr1+/Cspg4+ cells began at 2-4 days, significantly increased during the first 7 days, and stabilized toward the end of a 28-day implantation period (**Fig. 6b**, *p* < 0.01, one-way ANOVA with Tukey’s HSD test). During this time, the number of CX3cr1+/Cspg4+ cells significantly increased in very close spatial proximity (<50 μm) to the implanted electrode (**Fig. 6c**, *p* < 0.05, one-way ANOVA with Tukey’s HSD test). Investigation of the cortical depth in which these cells were observed revealed their positioning within 0-200 μm below the surface of the brain (layer I-II) (**Fig. 6d**). CX3cr1+/Cspg4+ cells demonstrated active extension and retraction of processes toward the surface of the electrode over imaging short durations (<30 min) at any given time point (**Fig. 6e**). Finally, CX3cr1+/Cspg4+ cells were most motile within the first week of electrode implantation, before stabilizing over the electrode surface between 2-4 weeks post-implantation (**Fig. 6f**). Patterns of motility seen in our visual findings were similarly demonstrated in quantification of the motility of different cellular compartments of CX3cr1+/Cspg4+ cells, revealing high soma movement within the first week before decreasing over the following 2-3 weeks post-implantation (**Fig. 6g-h**). Our results reveal for the first time a specialized population of reactive immune cells to the chronic implantation of a microelectrode array.

**Figure 6.**
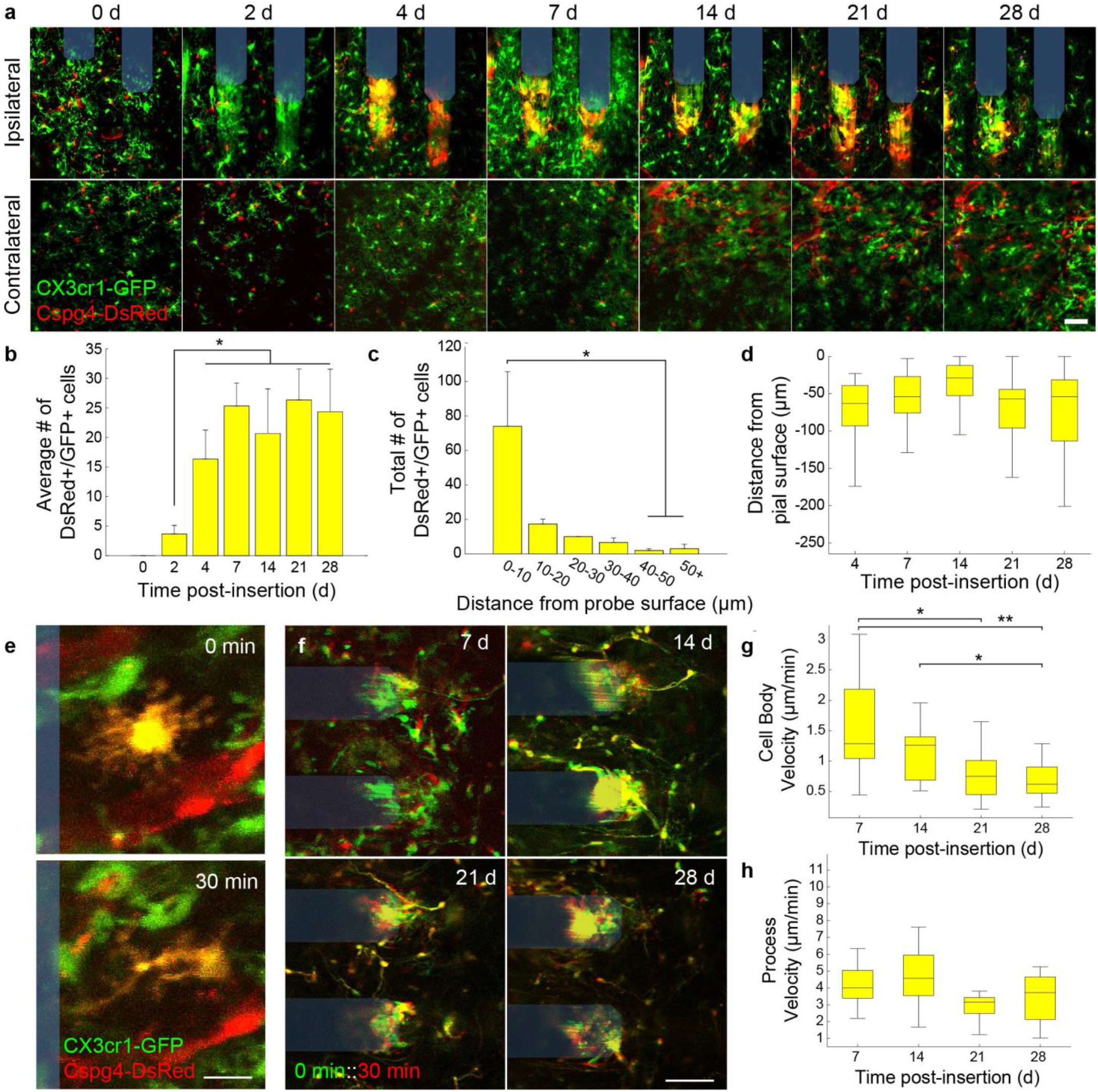
Emergence of reactive dual-CX3cr1+/Cspg4+ expressing cell population around chronically implanted microelectrodes. (a) Two-photon representative images of implanted microelectrodes (shaded blue) and contralateral control region over 28 day implantation period in a Cspg4-DsRed::CX3cr1-GFP mouse. Scale bar = 50 μm. (b) Average number of DsRed+/GFP+ cells counted over time post-insertion around chronically implanted microelectrode arrays. (c) Total number of DsRed+/GFP+ cells counted with respect to spatially binned distance away from the microelectrode surface. (d) Average distance of DsRed+/GFP+ with respect to pial surface over a 28 day implantation period. (e) Representative two-photon image demonstrating transition of DsRed+/GFP+ cell from a ramified to activated state over a 30 minute period. Scale bar = 10 μm (f) Superimposition of two time intervals 30 min apart demonstrating the change in motility of DsRed+/GFP+ cells over a 28 day implantation period. Note: Red vs. green labeling distinguishes between earlier (0 min, *green*) vs. later (30 min, *red*) timepoints, highlighting stationary features in yellow. Scale bar = 50 μm. (g) Average cell body velocity of DsRed+/GFP+ cells over a 28 day implantation period. (h) Average process velocity of DsRed+/GFP+ cells over a 28 day implantation period. *N* = 4 mice. * *p* < 0.05, ** *p* < 0.01. All data is reported as mean ± SEM.

### 4.6. Intracortical microstimulation modulates pericyte calcium dynamics in an amplitude- and frequency-dependent manner

The interplay between neurons and pericytes forms the basis behind neurovascular coupling. To understand how different patterns of neural activity can modulate pericyte Ca^2+^, we implanted a stimulating microelectrode in *PDGFRβ-GCaMP6s* mice (**Fig. 7a**). Electrical stimulation was administered using cathodic-leading, biphasic, charge-balanced waveforms with varying stimulating amplitudes (5, 10, and 15 µA) and frequencies (10 Hz burst, 10 Hz, 40 Hz, and 100 Hz) delivered over a 30-second stimulation duration (**Fig. 7b**). Modulating neural activity through intracortical microstimulation resulted in a visible decrease in pericyte calcium during the stimulation period before returning back to pre-stimulation levels (**Fig. 7c**). We determined that the amplitude of electrical stimulation can modulate the calcium activity of sphincter pericytes, highly contractile cells at the surface of the brain, as well as the diameter of the underlying vasculature, with 15 µA eliciting the largest change in magnitude compared to 5 and 10 µA (**Supplementary Fig. 1**). Therefore, we used 15 µA stimulating waveforms to determine whether there was any additional modulation based on varying frequency of electrical stimulation. We observed a clear modulatory effect of increasing frequency of stimulation on changes in pericyte calcium fluorescence (**Fig. 7d**). Specifically, we observed a significant decrease in mean ΔF/F_0_ at 40 Hz frequency of stimulation compared to 10 Hz burst or 10 Hz continuous stimulation throughout the stimulation period (**Fig. 7e**, *p* < 0.05 for 10 Hz burst vs. 40 Hz, *p* < 0.01 for 10 Hz vs. 40 Hz, one-way ANOVA with Tukey’s HSD test) as well as a significantly greater area over the ΔF/F_0_ curve at 40 and 100 Hz stimulation compared to 10 Hz burst or 10 Hz continuous (**Fig. 7g**, *p* < 0.05 for 10 Hz burst vs. 40 Hz, *p* < 0.01 for 10 Hz vs. 40 Hz, *p* < 0.05 for 10 Hz vs. 100 Hz, one-way ANOVA with Tukey’s HSD test). When looking at duration of pericyte Ca^2+^ activation during the electrical stimulation period, we observed an increase in response duration at 40 Hz stimulation compared to all other stimulation frequencies but did not detect any significant differences (**Fig. 7f**, *p* > 0.05, one-way ANOVA with Tukey’s HSD test). We discovered similar patterns of frequency modulation on intracellular Ca^2+^ dynamics within sphincter pericytes following intracortical microstimulation but with a greater change in magnitude (**Supplementary Fig. 2a-g**). We could not determine whether there were any changes in capillary tone alongside changes in calcium activity of adjacent pericytes during stimulation (*data not shown*), most likely due to the resolution constraints of our *in vivo* imaging system. However, we were able to detect a modulatory effect of stimulation frequency on the diameter of large arteriole blood vessels near sphincter pericytes (**Supplementary Fig. 2h**). Similar to the changes in intracellular pericyte calcium, increasing stimulation frequency resulted in large magnitude increases in diameter of underlying blood vessels (**Supplementary Fig. 2i-k**). Taken together, these results demonstrate that pericyte calcium and, by extension, vascular function can be modulated by varying the strength and activation patterns of local neural activity.

**Figure 7.**
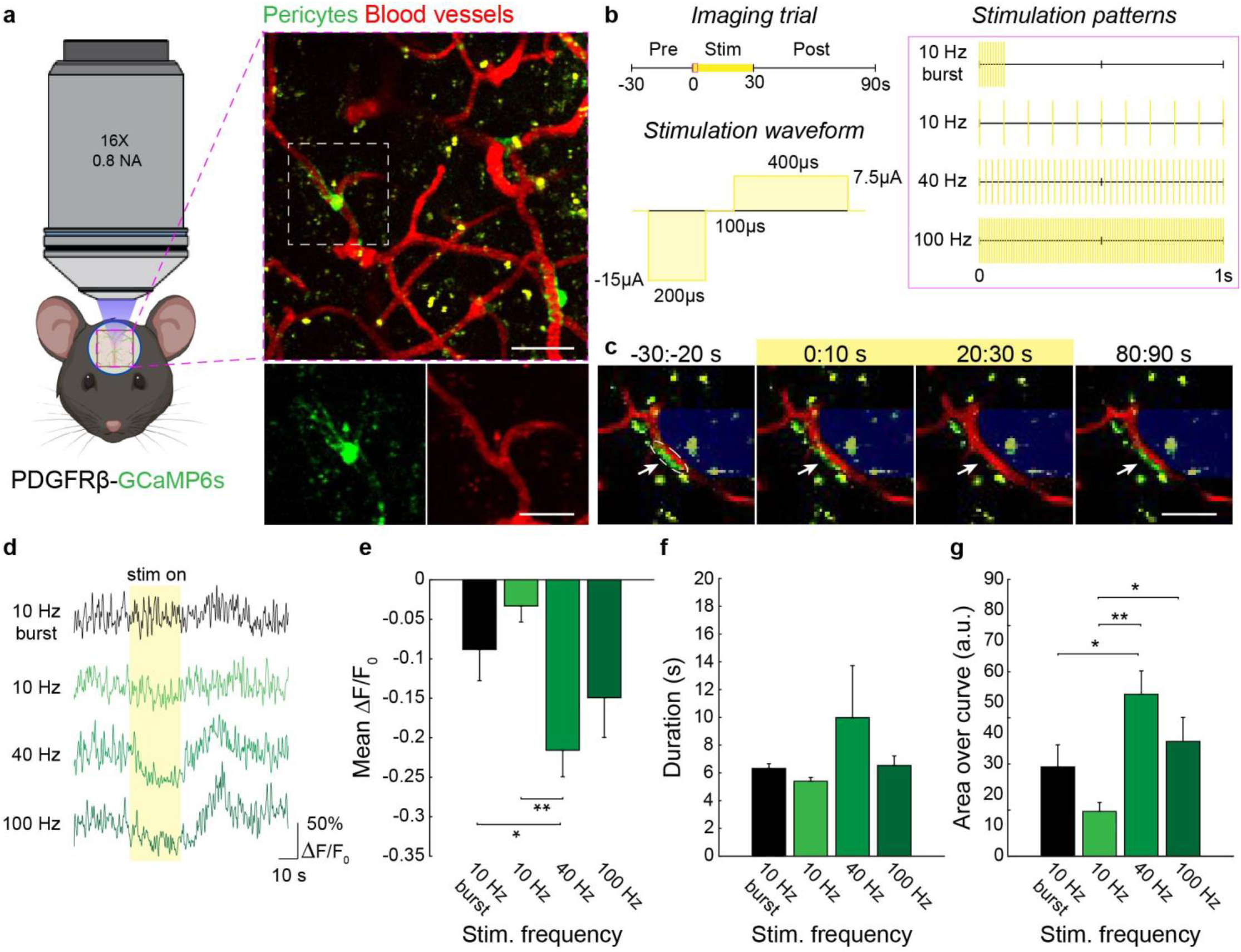
Intracortical microstimulation modulates pericyte Ca^2+^ in a frequency-dependent manner. (a) Two-photon visualization of pericyte Ca^2+^ dynamics during intracortical microstimulation (*electrode not shown*) using PDGFRβ-GCaMP6s mice. Scale bars = 10 µm. (b) Intracortical microstimulation waveforms consisted of cathodic-leading, biphasic, charge-balanced electrical pulses delivered at 10 Hz burst, 10 Hz, 40 Hz, or 100 Hz frequencies over a 30 second stimulation period. (c) Example of decrease in Ca^2+^ of capillary pericyte (*green*) adjacent to the stimulating electrode *(shaded blue*). Underlying blood vessels were visualized using SR101 (*red*). Scale bar = 10 µm. (d) Example Ca^2+^ traces during intracortical microstimulation at 10 Hz burst, 10 Hz, 40 Hz, or 100 Hz frequency. (e-g) Mean ΔF/F_0_, duration, and area over the curve during stimulation period at 10 Hz burst, 10 Hz, 40 Hz, or 100 Hz frequency (*N* = 3 mice, *n* = 5 cells). * *p* < 0.05, ** *p* < 0.01. All data is reported as mean ± SEM.

## 5.0. DISCUSSION

Penetrating microelectrodes allow researchers to interface directly with the brain and manipulate discrete brain cell populations in order to understand the function and dysfunction of our nervous system. While the long-term application of recording or stimulating electrodes can be used to diagnose and treat neurological brain dysfunction, their performance longevity and fidelity are limited by a severe foreign body response to an indwelling object within the brain. Efforts to identify the biological culprits responsible for a gradual decline in electrode performance have been obstructed by variable outcomes in the brain’s response to implanted probes. One potential source of that variability may be due to a yet poorly understood impact of chronic electrode implantation on perivascular cells within the brain. Pericytes are brain mural cells responsible for governing a variety of neuroimmune, homeostatic, and regulatory functions related to neuronal network activity and neurovascular coupling [19, 40-46]. Abnormal pericyte pathology has been emerging more as a key hallmark in the neurodegenerative sequence of events of a variety of brain injury conditions as well as disease [21-23,38,44,47-51]. Using intravital imaging, we assess the impact of chronic microelectrode implantation on the structure and function pericytes within the mouse cortex. Our results reveal distinct phases of both acute and chronic responses of pericytes to neural electrode implantation. Using dual-reporting fluorescent transgenic models. We report for the first time the discovery of a novel subset of reactive immune cells around implanted electrodes from our *in vivo* investigations. Finally, using electrical microstimulation we demonstrate that pericyte and vascular activity can be modulated in an amplitude- and frequency-dependent manner.

### 5.1. Acute pericyte dysfunction in response to microelectrode implantation

We revealed that insertion of a microelectrode within the brain immediately increases the intracellular Ca^2+^ levels of nearby pericytes (Fig. 2). Neurons have also shown to transiently increase their intracellular Ca^2+^ activity following microelectrode implantation [30]. One possibility for the sequence of events observed here is that these activated pericytes are responding to electrical or chemical signals emitted from activated neurons following electrode insertion and thereby elevating their own intracellular Ca^2+^ levels [52]. Previous work has shown neurotransmitters, including dopamine, noradrenealine, and ATP, have direct effects on pericytes, leading to the contraction of pericyte-associated vasculature [44, 53, 54]. Intracellular Ca^2+^ activity within pericytes is invariably tied to their contractile function. However, due to the limitations in laser scan settings and image resolution, we were unable to discern whether activated pericytes during electrode insertion induced constriction of underlying capillary vessels. Nevertheless, the strength of Ca^2+^ responses was observed to be more pronounced in pericyte soma compared to their processes (Fig. 2e-f). This is not surprising considering the circumferential processes responsible for wrapping around capillaries and mechanically contracting the endothelial lumen are positioned closest to pericyte soma [45]. In agreement with this idea, we demonstrate that pericytes produce maximal constriction of capillary vessels at the position of their cell soma (Fig. 2c). Similar results have been demonstrated in the presence of amyloid β oligomers within a rodent model of Alzheimer’s disease, which act on pericytes by inducing pericyte constriction of microvessels [48]. We also similarly observed obstructions in capillary flow at the site of pericyte soma positioned nearby an implanted microelectrode, suggesting their contractility to electrode insertion can contribute to reductions in cerebral blood flow (Fig. 2d). While the concept of constricted microvessels and stalled blood flow around indwelling electrodes has been proposed previously [7, 31], this is the first report to our knowledge implicating pericytes in the neurovascular dysfunction incurred during electrode implantation.

### 5.2. Pericytes proliferate and facilitate new vessel formation during the chronic response to electrode implantation

We previously revealed a spatiotemporal pattern of glial proliferation around chronically implanted microelectrodes [12]. In this same study, we also reported on an increased number of PDGFRβ+ and NG2+ pericytes at 7 days post-implantation of a microelectrode array. Here, we show that this increase in number of pericytes is due to cellular proliferation within the first 7- and 28-days post-implantation (Fig. 3). Similar time courses of pericyte proliferation and influx into damaged tissue regions following stroke have been reported [23]. During this time of elevated cellular proliferation, we observe pericytes facilitate angiogenesis around chronically implanted microelectrodes (Fig. 4). Angiogenesis is a function unique to pericytes and is triggered by the binding of platelet-derived growth factor BB (PDGF-BB) secreted from sprouting endothelial tip cells onto platelet-derived growth factor receptor β (PDGFR-β) expressed on pericyte membranes [55]. This endothelial cell signaling recruits pericytes to areas required for re-vascularization and allows pericytes to establish contact with sprouting vessels, directing and stabilizing the endothelial growth. Angiogenesis is a regenerative tissue response following conditions of tissue hypoxia, such as in stroke, or in the case of high cellular metabolic demands, such as within the tumor microenvironment. The potential for of ischemic injury following microelectrode implantation has been reported before through hemodynamic imaging of intrinsic signals around a chronically implanted microelectrode [13]. Pericytes are also known to facilitate leukocyte infiltration following severe brain injury. The generation of new blood vessels around chronically implanted electrodes could be a means to recruit peripheral immune cells to the electrode implant. Indeed, infiltration macrophages has previously been shown to consist of a significant population of immune cells around an implanted microelectrode array [56]. However, whether this presents a detriment to the survival of neural tissue around chronically implanted microelectrodes remains to be explored.

### 5.3. Appearance of a distinct subset of reactive immune cells around chronically implanted microelectrodes

To our surprise, we witnessed the emergence of a phenotypically distinct population of reactive cells within the electrode-tissue interface through the generation of a dual-fluorescent transgenic mouse model. These cells expressed both *Cspg4* and *Cx3cr1* promoters given their simultaneous 1:1 expression of DsRed and GFP fluorophores, respectively. No dual-labeled Cspg4+/CX3cr1+ cells were seen within the contralateral hemisphere, suggesting they appear specifically in response to the injury caused by electrode insertion (Fig. 6a). Cspg4+/CX3cr1+ cells first appeared 2-4 days after implantation and their numbers peaked by the end of the first week (Fig. 6b). The distribution of these cells was spatially confined to tissue areas immediately adjacent to the electrode surface (<50 µm). To our knowledge, this is the first report of its kind characterizing the appearance and behavior of a novel population of reactive cells within the electrode-tissue interface. One possibility is that these cells are a reactive subset of pericytes (i.e. *Cspg4*-expressing cells upregulating expression of *CX3cr1* promoter) operating within the fibrotic scar region of chronically implanted microelectrodes. Previous studies have reported on the ability for pericytes to develop microglia-like phenotypes in conditions of severe brain injury, such as stroke [27, 28]. In contrast, others have reported no evidence of pericyte expression of microglial markers by pericytes in other pathological conditions, such as acute brain injury [57]. This may be explained by differences in pathological conditions in which these cells were observed. Alternatively, these cells could constitute a subset of reactive in response to electrode implantation injury microglia (i.e. CX3cr1-expressing cells upregulating expression of *Cspg4* promoter). This possibility is more plausible given that microglia have demonstrated previously to upregulate the NG2 proteoglycan acutely in response to transient brain injury [57]. It is not clear from the data acquired here whether these injury-responsive cells are fundamentally neuroprotective or neurotoxic to the tissue microenvironment around chronic electrodes. Transcriptomic analyses of populations within mouse models of Parkinson’s and Alzheimer’s disease determined that CX3cr1+ microglia upregulate *Cspg4* expression specifically in the context of neurodegenerative conditions [58]. These *Cspg*^*high*^ microglial cells were found to be highly proliferative, due to upregulation of genes associated with the cell cycle, and differ functionally from disease-associated microglia (DAM), which are more phagocytic in nature [58]. Furthermore, transitional morphologies and patterns of cell motility and behavior of the cells characterized in this paper are reminiscent of other tissue scavengers commonly observed near the electrode surface, such as activated microglia and macrophages [31, 33]. Future work should focus on characterizing the functional role of this reactive population of immune cells around chronically implanted microelectrodes.

### 5.4 Modulation of neurovascular function using intracortical microstimulation

We have previously shown that neuronal activity can be modulated electrically by differing stimulation parameters, such as amplitude, frequency, or temporal patterning [59-63]. Moreover, different stimulation parameters cause varying degree of metabolic activity [62], and greater metabolic activity during stimulation is related to sustained neuronal depression [63]. Here, we show that, similar to neurons, the calcium activity within pericytes can be modulated by varying different stimulation parameters (Fig. 7). We report both decreases in intracellular calcium and proportional increases in underlying vascular tone when increasing the amplitude, or strength, of electrical stimulation (Fig. 7 and Supplementary Fig. 1 and 2). These effects on calcium fluorescence and vessel diameter were more readily observed in superficial sphincter cells, which populate larger arteriole blood vessels closer to the surface of the cortex [64], than in downstream capillary pericytes on higher-order branches of the vascular tree. Considering that the contractile proteins within mural cells necessary to exert mechanical changes in vascular tone are calcium-dependent, it is not surprising that larger changes in pericyte Ca^2+^ are accompanied by similar changes in vessel diameter. When varying stimulation frequency, we determined that higher frequencies of electrical stimulation such as 40 and 100 Hz produced greater decreases in mean ΔF/F and duration of activation compared to a lower 10 Hz frequency (Fig. 7d-g). We did not detect any difference in mean ΔF/F or duration of activation when varying temporal pattern of stimulation (10 Hz burst versus 10 Hz continuous).

One possible explanation of our results could be due to direct activation of pericytes by electrical stimulation. Pericytes express voltage-sensitive ion channels [45], suggesting their membrane potentials can be directly influenced by an applied electric field [65]. Alternatively, our findings could be the result of an indirect activation of pericyte function by stimulating neuronal activity. One possibility is that higher stimulation frequencies recruit different sub-populations of neurons (i.e. excitatory versus inhibitory neurons) compared to lower frequencies [60]. We have shown previously that increasing stimulation frequency (>100 Hz) recruits more sparse populations of neuronal cell bodies and results in an overall decrease in neuronal calcium activity when compared to lower frequencies of stimulation (<10 Hz). Inhibitory neurons, which are sparsely populated, are known to release vasoactive compounds and therefore can directly modulate neurovascular activity [66]. Our findings could therefore be the result of an increase in recruitment of inhibitory populations due to varying the frequency of stimulation [59, 61]. Furthermore, specific populations of inhibitory neurons, such as somatostatin-or nitric oxide synthase-expressing cells, have a more profound impact on cerebral blood flow regulation than others [67]. Future work should focus on improving our understanding of the mechanisms underlying the recruitment of various neuronal and non-neuronal populations during intracortical microstimulation as well as exploring how different patterns of electrical stimulation can be leveraged to modulate neurovascular function.

### 5.5 Limitations of study & future directions

Despite being identified over a century ago, it is only within the last couple of decades that the physiology and pathology of pericytes were beginning to be understood [43, 68]. This could be partly because pericytes are increasingly emerging as a heterogeneous group of vascular cells within the brain with a wide spectrum of genetic, structural, morphological, anatomical, physiological, and neuropathological profiles [36, 69]. In the past, an inability for researchers to agree on how to robustly categorize these cells properly across different experimental studies has limited the translational application of new knowledge [43]. For example, unlike neurons, microglia, and astrocytes, there is not one biomarker currently known that identifies only perivascular pericytes. Common pericyte markers such as NG2 or PDGFRβ are shared in some capacity by other structurally and functionally distinct cell populations within the brain. Therefore, accurate identification and characterization of pericyte cells depends on the co-localization of two or more markers to rule out other cell populations. Furthermore, a severe lack of appropriate investigative tools and experiments models has added to the difficulty in effective study of brain pericytes in the past.

The recent development of more neuroscientific techniques and advanced animal models have shown promise on revealing the distinct role of pericytes in brain function and dysfunction [70]. Some in vivo dyes were found to only be taken up by certain cell populations within the brain, such as NeuroTrace 500/525 in pericytes [71], and have demonstrated some success in the specific investigation of capillary pericytes. However, this dye requires topical application onto the brain and is therefore only applicable in studies which utilize an open skull preparation. In our study, we use an *Cspg4-DsRed* mouse model which constitutively expresses a red fluorescent protein within pericyte cells in the brain [72]. This has been commonly used to assess pericyte physiology and pathology within the brain in the past, which can be accurately performed by identifying and characterizing cells which demonstrate the same anatomical, morphological, and structural properties as classically identified brain pericytes (i.e. anatomical position along blood vessels with “bump on a log” morphology) [44, 64, 73]. However, in certain pathological conditions which promote mural cell reactivity and pericytosis, such as detachment from blood vessels and migration into the scarred region following brain injury, the assessment of pericyte fate and function becomes more difficult.

The *Cspg4* gene encodes for the NG2 proteoglycan, which is not only expressed in pericytes but also known to be commonly expressed within oligodendrocyte precursor cells (NG2 glia), as well as microglia and astrocytes in certain pathological conditions [57]. We previously characterized the spatiotemporal reactivity of NG2 glia around implanted microelectrodes [33]. Given the limitation in the model used, it is difficult to say whether the DsRed+ cells we observed encapsulating the electrode surface are partly from DsRed+ pericyte cells or if they consist of other glial cells which upregulate expression of NG2 following implantation injury. One way to discern whether the cells we observe within this scarred region are pericytes or not would be to employ more advance transgenic mouse lines which express fluorophores under multiple pericyte promoters, such as the NG2 and PDGFRβ promoters in a recently developed pericyte-CreER line [47]. Alternatively, use of single cell genetic sequencing to better understand gene expression profiles will offer profound insight on the origin, fate, and function of these seemingly phenotypically-similar reactive cell populations interacting at the electrode-tissue interface [58].

## 6.0. CONCLUSION

Development of advanced neural electrode technology is accelerating our understanding of nervous system health and disease at a rapid pace. Yet, despite their potential as tools for neuroscience discovery and clinical rehabilitation, intracortical electrodes regularly suffer from depreciations in detecting and modulating brain activity as a result of severe biological responses to a foreign body. Due to the inherent complexity of the brain’s immune response to device injury, the mechanisms which govern biological failure modes of neural implants remain unknown. Efforts to pinpoint exact biological correlates of device performance are confounded by a poorly understood heterogeneity of immune cells (i.e. myeloid cells) within the brain which are known regulate tissue homeostasis, inflammation, and wound repair following injury. Brain pericytes are one such myeloid population whose contribution to the bodily tissue reactions to implanted electrodes is currently unknown. Here, we employed two-photon imaging of multiple transgenic mouse models over a 28-day implantation period to elucidate dynamic changes in the fate and behavior of pericyte cells during the course of microelectrode implantation. We discovered that pericytes are dynamic in the presence of chronically implanted electrodes, initially constricting around capillary microvessels in response to electrode insertion before actively engaging in tissue remodeling at the electrode-tissue interface. During our investigation we also reveal distinct population reactive immune cells present within the scar region of chronically implanted microelectrodes. Finally, we established that the calcium activity of pericytes can be altered through intracortical microstimulation in an amplitude- and frequency-dependent manner. Future work further characterizing the heterogeneity in brain tissue responses of the pericyte population and their pathological roles within the brain will facilitate the development of more biologically adaptive implants and innovative therapeutic strategies to combat the body’s inherent immune response to brain injury and disease.

## 7.0. ACKNOWLEDGEMENTS

This work was supported by NIH NINDS R21NS108098, NIH NINDS R01NS094396 and a diversity supplement to the parent grant, NIH NINDS R01NS105691, NIH NINDS R01NS115707 and a diversity supplement to the parent grant, R01NS129632, NSF CBET CAREER 1943906, and NIH NINDS F99NS124186.

---

**Supplementary Figure 1.**
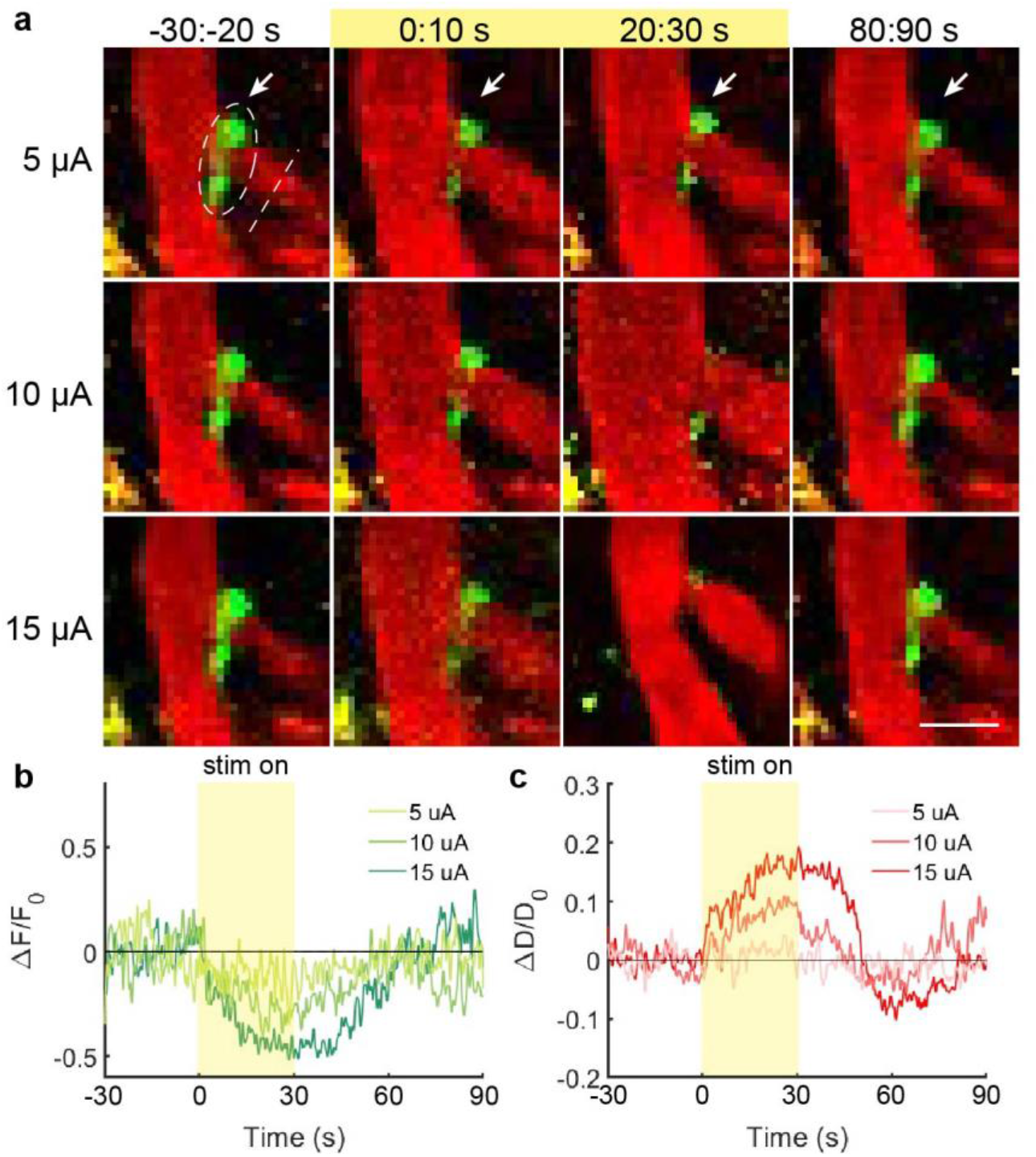
Intracortical microstimulation modulates sphincter pericyte Ca^2+^ and vessel diameter in an amplitude-dependent manner. (a) Example of decrease in Ca^2+^ of sphincter pericyte (*green*) and increase in diameter of underlying vasculature (*red*) during stimulation period at 5, 10, and 15 µA stimulation amplitudes. Scale bar = 10 µm. (b) Intracortical microstimulation results in a decrease in sphincter pericyte Ca^2+^ in an amplitude-dependent manner. (c) Intracortical microstimulation results in an increase in diameter of underlying vasculature in an amplitude-dependent manner.

**Supplementary Figure 2.**
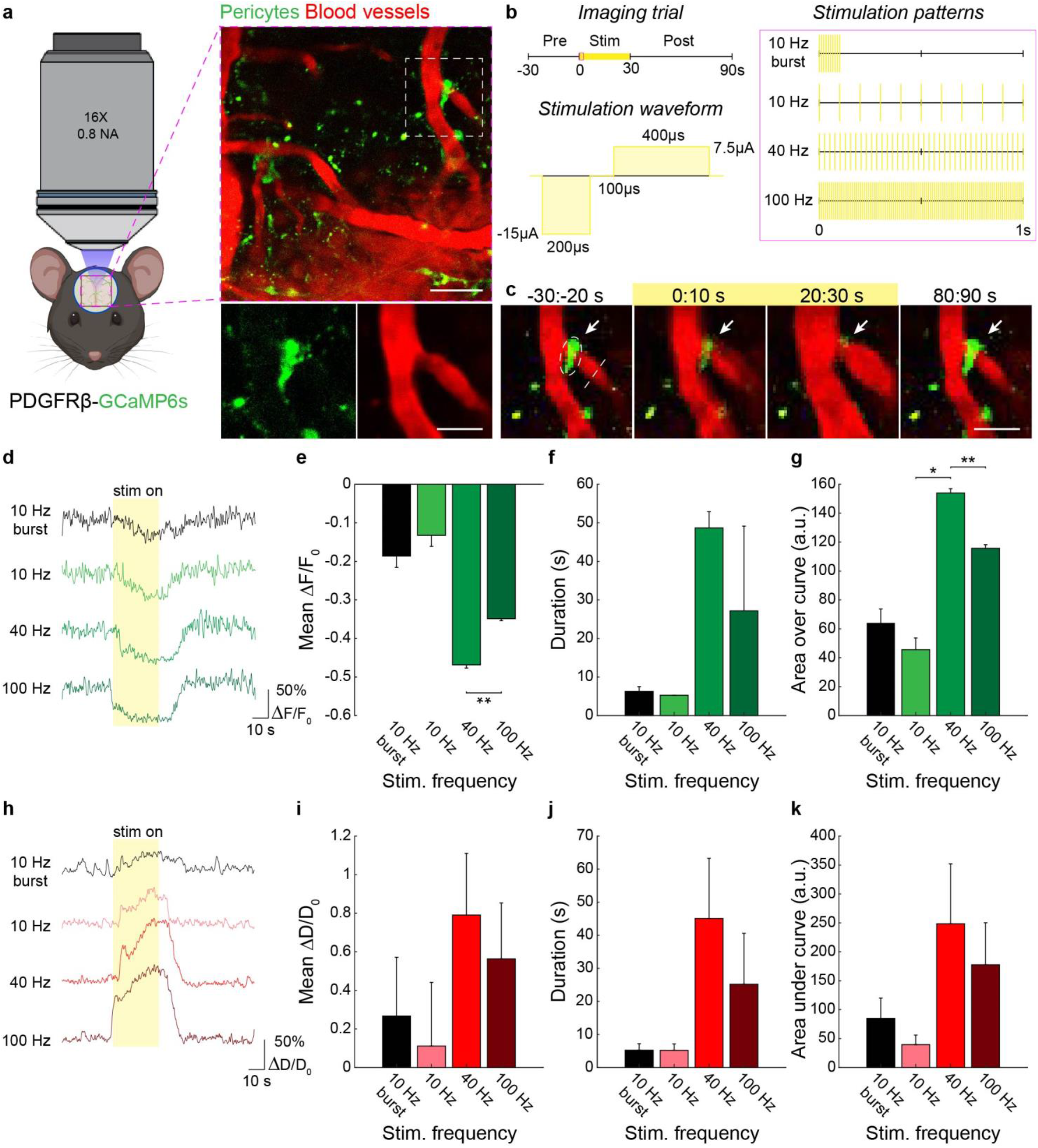
Intracortical microstimulation modulates sphincter pericyte Ca^2+^ and vessel diameter in a frequency-dependent manner. (a) Two-photon visualization of sphincter pericyte Ca^2+^ dynamics and underlying vascular tone during intracortical microstimulation (*electrode not shown*) using PDGFRβ-GCaMP6s mice. Scale bars = 10 µm. (b) Intracortical microstimulation waveforms consisted of cathodic-leading, biphasic, charge-balanced electrical pulses delivered at 10 Hz burst, 10 Hz, 40 Hz, or 100 Hz frequencies over a 30 second stimulation period. (c) Example of decrease in Ca^2+^ of sphincter pericyte (*green*) and increase in diameter of underlying vasculature (*red*) during stimulation period. Scale bar = 10 µm. (d) Example Ca^2+^ traces during intracortical microstimulation at 10 Hz burst, 10 Hz, 40 Hz, or 100 Hz frequency. (e-g) Mean ΔF/F_0_, duration, and area over the curve during stimulation period at 10 Hz burst, 10 Hz, 40 Hz, or 100 Hz frequency. (h) Example diameter traces during intracortical microstimulation at 10 Hz burst, 10 Hz, 40 Hz, or 100 Hz frequency. (i-k) Mean ΔD/D_0_, duration, and area under the curve during stimulation period at 10 Hz burst, 10 Hz, 40 Hz, or 100 Hz frequency. * *p* < 0.05, ** *p* < 0.01. All data is reported as mean ± SEM.

## REFERENCES

1. Steinmetz, N.A., et al., Neuropixels 2.0: A miniaturized high-density probe for stable, long-term brain recordings. Science, 2021. 372(6539).

2. Lee, K., et al., Flexible, scalable, high channel count stereo-electrode for recording in the human brain. Nature communications, 2024. 15(1): p. 218.

3. Lycke, R., et al., Low-threshold, high-resolution, chronically stable intracortical microstimulation by ultraflexible electrodes. Cell reports, 2023. 42(6).

4. Pérez-Prieto, N. and M. Delgado-Restituto, Recording strategies for high channel count, densely spaced microelectrode arrays. Frontiers in Neuroscience, 2021. 15: p. 681085.

5. Hughes, C.L., et al., Perception of microstimulation frequency in human somatosensory cortex. Elife, 2021. 10: p. e65128.

6. Flesher, S.N., et al., Intracortical microstimulation of human somatosensory cortex. Science Translational Medicine, 2016. 8(361): p. 361ra141–361ra141.

7. Kozai, T.D., et al., Brain tissue responses to neural implants impact signal sensitivity and intervention strategies. ACS chemical neuroscience, 2015. 6(1): p. 48–67.

8. Szymanski, L.J., et al., Neuropathological effects of chronically implanted, intracortical microelectrodes in a tetraplegic patient. Journal of Neural Engineering, 2021. 18(4): p. 0460b9.

9. Urdaneta, M.E., et al., The long-term stability of intracortical microstimulation and the foreign body response are layer dependent. Frontiers in Neuroscience, 2022. 16: p. 908858.

10. Wellman, S.M., F. Cambi, and T.D. Kozai, The role of oligodendrocytes and their progenitors on neural interface technology: a novel perspective on tissue regeneration and repair. Biomaterials, 2018. 183: p. 200–217.

11. Wellman, S.M. and T.D. Kozai, Understanding the inflammatory tissue reaction to brain implants to improve neurochemical sensing performance. 2017, ACS Publications. p. 2578–2582.

12. Wellman, S.M., et al., Revealing spatial and temporal patterns of cell death, glial proliferation, and blood-brain barrier dysfunction around implanted intracortical neural interfaces. Frontiers in neuroscience, 2019. 13: p. 493.

13. Michelson, N.J., et al., Multi-scale, multi-modal analysis uncovers complex relationship at the brain tissue-implant neural interface: new emphasis on the biological interface. Journal of neural engineering, 2018. 15(3): p. 033001.

14. Rousche, P.J. and R.A. Normann, Chronic recording capability of the Utah Intracortical Electrode Array in cat sensory cortex. Journal of neuroscience methods, 1998. 82(1): p. 1–15.

15. Williams, J.C., R.L. Rennaker, and D.R. Kipke, Long-term neural recording characteristics of wire microelectrode arrays implanted in cerebral cortex. Brain Research Protocols, 1999. 4(3): p. 303–313.

16. Kozai, T.D.Y., et al., Reduction of neurovascular damage resulting from microelectrode insertion into the cerebral cortex using in vivo two-photon mapping. Journal of neural engineering, 2010. 7(4): p. 046011.

17. Bjornsson, C., et al., Effects of insertion conditions on tissue strain and vascular damage during neuroprosthetic device insertion. Journal of neural engineering, 2006. 3(3): p. 196.

18. Sweeney, M.D., S. Ayyadurai, and B.V. Zlokovic, Pericytes of the neurovascular unit: key functions and signaling pathways. Nature neuroscience, 2016. 19(6): p. 771–783.

19. Kozberg, M. and E. Hillman, Neurovascular coupling and energy metabolism in the developing brain. Progress in brain research, 2016. 225: p. 213–242.

20. Kozai, T.D., et al., Effects of caspase-1 knockout on chronic neural recording quality and longevity: insight into cellular and molecular mechanisms of the reactive tissue response. Biomaterials, 2014. 35(36): p. 9620–9634.

21. Dalkara, T., L. Alarcon-Martinez, and M. Yemisci, Pericytes in ischemic stroke, in Pericyte Biology in Disease. 2019, Springer. p. 189–213.

22. Di, Z., et al., Effect of pericytes on cerebral microvasculature at different time points of stroke. BioMed Research International, 2021. 2021.

23. Fernández-Klett, F., et al., Early loss of pericytes and perivascular stromal cell-induced scar formation after stroke. J Cereb Blood Flow Metab, 2013. 33(3): p. 428–39.

24. Hess, D.C., et al., Hematopoietic origin of microglial and perivascular cells in brain. Experimental neurology, 2004. 186(2): p. 134–144.

25. Ozerdem, U. and W.B. Stallcup, Early contribution of pericytes to angiogenic sprouting and tube formation. Angiogenesis, 2003. 6(3): p. 241–249.

26. Halliday, M.R., et al., Accelerated pericyte degeneration and blood-brain barrier breakdown in apolipoprotein E4 carriers with Alzheimer’s disease. J Cereb Blood Flow Metab, 2016. 36(1): p. 216–27.

27. Sakuma, R., et al., Brain pericytes serve as microglia-generating multipotent vascular stem cells following ischemic stroke. Journal of neuroinflammation, 2016. 13(1): p. 1–13.

28. Özen, I., et al., Brain pericytes acquire a microglial phenotype after stroke. Acta neuropathologica, 2014. 128(3): p. 381–396.

29. Chen, K., et al., In vivo spatiotemporal patterns of oligodendrocyte and myelin damage at the neural electrode interface. Biomaterials, 2021. 268: p. 120526.

30. Eles, J.R., et al., In vivo imaging of neuronal calcium during electrode implantation: spatial and temporal mapping of damage and recovery. Biomaterials, 2018. 174: p. 79–94.

31. Kozai, T.D.Y., et al., In vivo two-photon microscopy reveals immediate microglial reaction to implantation of microelectrode through extension of processes. Journal of neural engineering, 2012. 9(6): p. 066001.

32. Savya, S.P., et al., In vivo spatiotemporal dynamics of astrocyte reactivity following neural electrode implantation. Biomaterials, 2022. 289: p. 121784.

33. Wellman, S.M. and T.D. Kozai, In vivo spatiotemporal dynamics of NG2 glia activity caused by neural electrode implantation. Biomaterials, 2018. 164: p. 121–133.

34. Kozai, T.D., et al., Two-photon imaging of chronically implanted neural electrodes: Sealing methods and new insights. Journal of neuroscience methods, 2016. 258: p. 46–55.

35. Barrett, M.J., et al., CHIPS: an extensible toolbox for cellular and hemodynamic two-photon image analysis. 2018, Springer.

36. McDowell, K.P., et al., VasoMetrics: unbiased spatiotemporal analysis of microvascular diameter in multi-photon imaging applications. Quantitative Imaging in Medicine and Surgery, 2021. 11(3): p. 969.

37. Nelson, A.R., et al., Channelrhodopsin excitation contracts brain pericytes and reduces blood flow in the aging mouse brain in vivo. Frontiers in aging neuroscience, 2020. 12: p. 108.

38. Costa, M.A., et al., Pericytes constrict blood vessels after myocardial ischemia. Journal of molecular and cellular cardiology, 2018. 116: p. 1–4.

39. Yemisci, M., et al., Pericyte contraction induced by oxidative-nitrative stress impairs capillary reflow despite successful opening of an occluded cerebral artery. Nature medicine, 2009. 15(9): p. 1031–1037.

40. Berthiaume, A.-A., et al., Dynamic remodeling of pericytes in vivo maintains capillary coverage in the adult mouse brain. Cell reports, 2018. 22(1): p. 8–16.

41. Berthiaume, A.-A., et al., Pericyte structural remodeling in cerebrovascular health and homeostasis. Frontiers in Aging Neuroscience, 2018. 10: p. 210.

42. Sá-Pereira, I., D. Brites, and M.A. Brito, Neurovascular unit: a focus on pericytes. Molecular neurobiology, 2012. 45: p. 327–347.

43. Attwell, D., et al., What is a pericyte? Journal of Cerebral Blood Flow & Metabolism, 2016. 36(2): p. 451–455.

44. Hall, C.N., et al., Capillary pericytes regulate cerebral blood flow in health and disease. Nature, 2014. 508(7494): p. 55–60.

45. Hamilton, N.B., D. Attwell, and C.N. Hall, Pericyte-mediated regulation of capillary diameter: a component of neurovascular coupling in health and disease. Front Neuroenergetics, 2010. 2.

46. Hartmann, D.A., et al., Brain capillary pericytes exert a substantial but slow influence on blood flow. Nature neuroscience, 2021. 24(5): p. 633–645.

47. Nikolakopoulou, A.M., et al., Pericyte loss leads to circulatory failure and pleiotrophin depletion causing neuron loss. Nature Neuroscience, 2019. 22(7): p. 1089–1098.

48. Nortley, R., et al., Amyloid β oligomers constrict human capillaries in Alzheimer’s disease via signaling to pericytes. Science, 2019. 365(6450): p. eaav9518.

49. Medina-Flores, F., et al., The active role of pericytes during neuroinflammation in the adult brain. Cellular and Molecular Neurobiology, 2023. 43(2): p. 525–541.

50. van Splunder, H., et al., Pericytes in the disease spotlight. Trends in Cell Biology, 2023.

51. Zhang, Y., et al., Pericytes in Alzheimer’s disease: Key players and therapeutic targets. Experimental Neurology, 2024: p. 114825.

52. Brown, L.S., et al., Pericytes and Neurovascular Function in the Healthy and Diseased Brain. Front Cell Neurosci, 2019. 13: p. 282.

53. Peppiatt, C.M., et al., Bidirectional control of CNS capillary diameter by pericytes. Nature, 2006. 443(7112): p. 700–704.

54. WU, D.M., et al., Dopamine activates ATP-sensitive K+ currents in rat retinal pericytes. Visual neuroscience, 2001. 18(6): p. 935–940.

55. Ribatti, D., B. Nico, and E. Crivellato, The role of pericytes in angiogenesis. Int J Dev Biol, 2011. 55(3): p. 261–8.

56. Ravikumar, M., et al., The roles of blood-derived macrophages and resident microglia in the neuroinflammatory response to implanted intracortical microelectrodes. Biomaterials, 2014. 35(28): p. 8049–64.

57. Huang, W., et al., Acute brain injuries trigger microglia as an additional source of the proteoglycan NG2. Acta neuropathologica communications, 2020. 8(1): p. 1–15.

58. Liu, Y.-j., et al., Cspg4high microglia contribute to microgliosis during neurodegeneration. Proceedings of the National Academy of Sciences, 2023. 120(8): p. e2210643120.

59. Eles, J.R., K.C. Stieger, and T.D. Kozai, The temporal pattern of intracortical microstimulation pulses elicits distinct temporal and spatial recruitment of cortical neuropil and neurons. Journal of neural engineering, 2021. 18(1): p. 015001.

60. Michelson, N.J., et al., Calcium activation of cortical neurons by continuous electrical stimulation: Frequency dependence, temporal fidelity, and activation density. Journal of neuroscience research, 2019. 97(5): p. 620–638.

61. Stieger, K.C., et al., Intracortical microstimulation pulse waveform and frequency recruits distinct spatiotemporal patterns of cortical neuron and neuropil activation. Journal of neural engineering, 2022. 19(2): p. 026024.

62. Suematsu, N., A.L. Vazquez, and T.D.Y. Kozai, Activation and depression of neural and hemodynamic responses induced by the intracortical microstimulation and visual stimulation in the mouse visual cortex. Journal of Neural Engineering, 2024.

63. Hughes, C. and T. Kozai, Dynamic amplitude modulation of microstimulation evokes biomimetic onset and offset transients and reduces depression of evoked calcium responses in sensory cortices. Brain Stimulation, 2023. 16(3): p. 939–965.

64. Zambach, S.A., et al., Precapillary sphincters and pericytes at first-order capillaries as key regulators for brain capillary perfusion. Proceedings of the National Academy of Sciences, 2021. 118(26): p. e2023749118.

65. Pulgar, V.M., Direct electric stimulation to increase cerebrovascular function. 2015, Frontiers Media SA. p. 54.

66. Lecrux, C. and E. Hamel, Neuronal networks and mediators of cortical neurovascular coupling responses in normal and altered brain states. Philosophical Transactions of the Royal Society B: Biological Sciences, 2016. 371(1705): p. 20150350.

67. Krawchuk, M.B., et al., Optogenetic assessment of VIP, PV, SOM and NOS inhibitory neuron activity and cerebral blood flow regulation in mouse somato-sensory cortex. Journal of Cerebral Blood Flow & Metabolism, 2020. 40(7): p. 1427–1440.

68. Rouget, C., Memoire sur le develloppment, la structure et les propietes physiologiques des capillaries senguins et lymphatiques. Arch. Physiol. Norm. Pathol., 1873. 5: p. 603–663.

69. Dias Moura Prazeres, P.H., et al., Pericytes are heterogeneous in their origin within the same tissue. Dev Biol, 2017. 427(1): p. 6–11.

70. Nielson, C.D. and A.Y. Shih, In vivo Single Cell Optical Ablation of Brain Pericytes. Front Neurosci, 2022. 16: p. 900761.

71. Damisah, E.C., et al., A fluoro-Nissl dye identifies pericytes as distinct vascular mural cells during in vivo brain imaging. Nat Neurosci, 2017. 20(7): p. 1023–1032.

72. Zhu, X., D.E. Bergles, and A. Nishiyama, NG2 cells generate both oligodendrocytes and gray matter astrocytes. Development, 2008. 135(1): p. 145–57.

73. Mishra, A., et al., Imaging pericytes and capillary diameter in brain slices and isolated retinae. Nat Protoc, 2014. 9(2): p. 323–36.

